# Cortistatin neurons in the prelimbic cortex regulate seizure susceptibility in female mice via BDNF–TrkB signaling

**DOI:** 10.64898/2026.01.28.702318

**Authors:** Aaron J. Salisbury, Lourdes Figueroa, Keri Martinowich, Michael S. Totty

## Abstract

The risk of developing psychiatric disorders, particularly stress-related disorders such as major depressive disorder (MDD) and post-traumatic stress disorder (PTSD), is increased threefold in patients with epilepsy. While this increased risk may arise as a consequence of living with epilepsy, shared neurobiological mechanisms, particularly dysregulation of GABAergic signaling, may also contribute. To investigate this link, we investigated the function of GABAergic neurons co-expressing the neuropeptide cortistatin (CST), which has anticonvulsant effects and is implicated in both MDD and PTSD. Targeting CST+ neurons in the prelimbic cortex (PrL), a rodent brain region that is functionally and anatomically similar to the human dorsal anterior cingulate cortex (dACC), we found that ablating CST+ neurons disrupts context-dependent fear renewal, causes spontaneous convulsive seizures, dramatically increases susceptibility to chemically-induced seizures, and increases anxiety-like phenotypes following stressors. We further show that repeated chemogenetic inhibition of CST+ neurons increases the rate of seizure kindling in female mice, and that disruption of brain derived neurotrophic factor signaling in CST+ neurons phenocopies the effects of acute inhibition. These data support the hypothesis that epilepsy and stress-related psychiatric disorders potentially share common neurobiological mechanisms, and that loss of CST+ neuron function may be a critical feature underlying fear dysregulation and cortical hyperexcitability.

**Highlights:**

- Loss of prelimbic CST+ neurons disrupts fear renewal, causes anxiety-like symptoms, and induces spontaneous seizures.
- Inhibition of prelimbic CST+ by chemogenetics or BDNF inhibition by TrkB increases seizure susceptibility in females only.
- CST+ neurons represent a subset of primarily SST and PVALB expressing GABAergic neurons.

## 1 Introduction

A growing body of evidence suggests that epilepsy and stress-related disorders share common underlying neurobiological mechanisms. Patients with post-traumatic stress disorder (PTSD) have a 3 fold higher risk of developing epilepsy compared to the general population (Chen et al., 2017), and epilepsy patients have a 3 fold increased risk for developing PTSD (Pepi et al., 2024). In addition, patients who experience trauma frequently have coincident onset of PTSD symptoms and epilepsy (Soncin et al., 2021). Perturbations to excitatory/inhibitory balance arising from dysfunction in GABAergic cell populations are implicated in all of these disorders (Sessolo et al., 2015; Fang et al., 2018; Page & Coutellier, 2019; Sheth et al., 2019). Under normal conditions, inhibitory signaling supports signal transduction by filtering noise (Heise et al., 2022; Zhang et al., 2025) modulating gain (Root et al., 2008; Stange et al., 2013; Fu et al., 2014), gating excitatory inputs (Katz, 2003) and preventing runaway excitation (Silberberg, 2008). Dysfunction of specific subtypes of GABAergic neurons, including those expressing somatostatin (SST)-, parvalbumin (PVALB)-, or vasointestinal peptide (VIP), is implicated in a wide variety of psychiatric and neurological disorders, including major depressive disorder (MDD), PTSD, and epilepsy (Fee et al., 2017; Regev-Tsur et al., 2020; Zhao et al., 2024). For example, epilepsy is associated with dysfunction in SST+ neurons (Yekhlef et al., 2015; Y. Wang et al., 2020; Zheng et al., 2023), and our recent study in postmortem human brain tissue revealed that *CORT*, the gene encoding the neuropeptide cortistatin (CST), which is often expressed in SST+ neurons, was one of the top downregulated transcripts in the dorsal anterior cingulate cortex (dACC) of individuals with either MDD or PTSD (Jaffe et al., 2022). The dACC, as well as the prelimbic cortex (PrL), an anatomically and functionally similar region in the rodent brain, regulate fear expression (Milad et al., 2007; Burgos-Robles et al., 2009), and fear associated cue discrimination (Dunsmoor & LaBar, 2012; Stujenske et al., 2022). Based on these findings and observed comorbidities between epilepsy, PTSD, and MDD (Chen et al., 2017), (Pepi et al., 2024), we hypothesized that dysregulation in CST+ neurons may represent a common neural mechanism underlying symptoms in PTSD and epilepsy.

The neuropeptide CST has received relatively little attention since its discovery nearly thirty years ago (de Lecea et al., 1997). CST is structurally similar to SST and binds with high affinity to all four major SST receptors (Fukusumi et al., 1997). However, unlike SST, CST is expressed in both SST+ and PVALB+ neurons and is found almost exclusively in the cortex and cortical-like brain regions, including the hippocampus and amygdala. This is in contrast to localization of the broader SST+ and PVALB+ populations, which are found throughout the central nervous system. CST regulates sleep and antagonizes inflammation (de Lecea et al., 1996; Gonzalez-Rey et al., 2006; Castillo-González et al., 2024), and our lab previously found that constitutive ablation of CST+ neurons in early postnatal development causes fatal seizures in mice (J. L. Hill et al., 2019). CST is frequently co-expressed with other antiepileptic and anxiolytic neuropeptides, including SST, neuropeptide Y (NPY), and corticotropin-releasing hormone binding protein (CRHBP). These peptides are also implicated in stress-related disorders including MDD and PTSD (Enman et al., 2015; Fuchs et al., 2017), and similar to *CORT*, the transcripts encoding these peptides are downregulated in in the dACC in individuals with MDD and PTSD (Jaffe et al., 2022). Of note, transcription of these genes is regulated by brain-derived neurotrophic factor (BDNF) signaling, which plays key roles in the development, maturation, and physiological function of GABAergic neurons (Yamada et al., 2002; Hong et al., 2008; Gottmann et al., 2009). In line with these findings, our lab found that constitutive knockout of the BDNF receptor, TrkB, specifically in CST+ neurons, induces seizures in mice (J. L. Hill et al., 2019). Although this work establishes that BDNF-TrkB in CST+ neurons is critical for regulating excitability during development, whether CST+ neuron function is similarly required for restraining excitability in adult mice remains unknown.

To investigate the role of CST+ neurons in regulating hyperexcitability associated with fear, anxiety, and epilepsy, we assessed Pavlovian auditory fear conditioning, open-field locomotion, and seizure development in a pentylenetetrazol (PTZ) kindling model, after either regionally focused genetic ablation or acute inhibition of CST+ neurons in adult mice. Ablation of CST+ neurons in the PrL induced spontaneous seizures and impaired context-dependent fear renewal in both male and female mice, whereas chemogenetic inhibition and loss of BDNF-TrkB signaling in CST+ neurons in the PrL increased the rate of kindling exclusively in female mice. In summary, these data demonstrate a role for CST+ neurons in regulating fear and hyperexcitability in adult mice and demonstrate that BDNF-TrkB signaling in CST+ neurons is critical for preventing development of seizure activity in females.

## 2 Results

### 2.1 Ablation of CST+ neurons in the PrL increases fear renewal and seizure susceptibility

Because expression of *CORT* is dysregulated in post-traumatic stress disorder (PTSD) in the human dorsal anterior cingulate cortex (dACC) (Jaffe et al., 2022), a region that is implicated in fear expression (Milad et al., 2007), we hypothesized that loss of *Cort*-expressing (CST+) neurons may impair acquisition or extinction of fear learning. To test this hypothesis, we selectively ablated CST+ neurons in the mouse prelimbic cortex (PrL), a region that is functionally and anatomically similar to the human dACC, using a caspase-3 virus. Mice expressing cre recombinase under control of the endogenous *Cort* promoter (*Cort*-T2A-cre^+/-^) were bilaterally injected with a cre dependent mCherry control vector (AAV5-Ef1a-DIO-mCherry; Control group) or a mixture of a cre recombinase dependent caspase 3 virus and the mCherry control virus (AAV5-Ef1a-casp3 and AAV5-Ef1a-DIO-mCherry; Ablation group) (Fig. 1A). Four weeks following injections, mice were tested on an auditory fear conditioning paradigm (Fig. 1B). There were no differences between groups in freezing behavior during conditioning (Fig. 1C, *F*_(1,17)_ = 0.1347, *p* = 0.7182), context extinction (Fig. 1D, *F*_(1,17)_ = 1.389, *p* = 0.2589), cued extinction (Fig. 1E, *F*_(1,17)_ = 1.795, *p* = 0.1979; Fig. 1G, *F*_(1,17)_ = 3.637, *p* = 0.0735), or extinction retrieval (F_(1,16)_ = 0.3640, p = 0.5547). CST+ neuron ablation did, however, increase fear renewal when tested in a novel context (Fig. 1G, *F*_(1,16)_ = 5.356, *p* = 0.0343). There were also no observed differences in conditioning, context, extinction, cued extinction, or extinction retrieval between groups when stratified by sex (Fig. S1). Over the course of the fear conditioning experiments, we noted that a subset of mice in the Ablation group developed spontaneous tonic-clonic seizures (Fig. 1H). As seizures or locomotion issues could confound freezing behavior, we quantified locomotion and anxiety-like behavior in an open field. While no differences in locomotion were observed (Fig. 1K, *t* = 0.5050, *p* = 0.6204), CST+ neuron ablation decreased time spent in the center of the open field (Fig. 1J, *t* = 4.067, *p* = 0.0009), evidence of increased anxiety-like behavior. Since spontaneous seizures were observed in a subset of mice in the Ablation group, we administered the chemo-convulsant pentylenetetrazole (PTZ, 35 mg/kg) to test if ablating CST+ neurons increased sensitivity to developing seizures. While 35 mg/kg is considered a subacute dose in wild type mice, it caused fatal seizures in all eight mice in the Ablation group while only inducing minor symptoms in Control mice (Fig. 1I, *t* = 13.39, *p* < 0.0001).

**Fig. 1.**
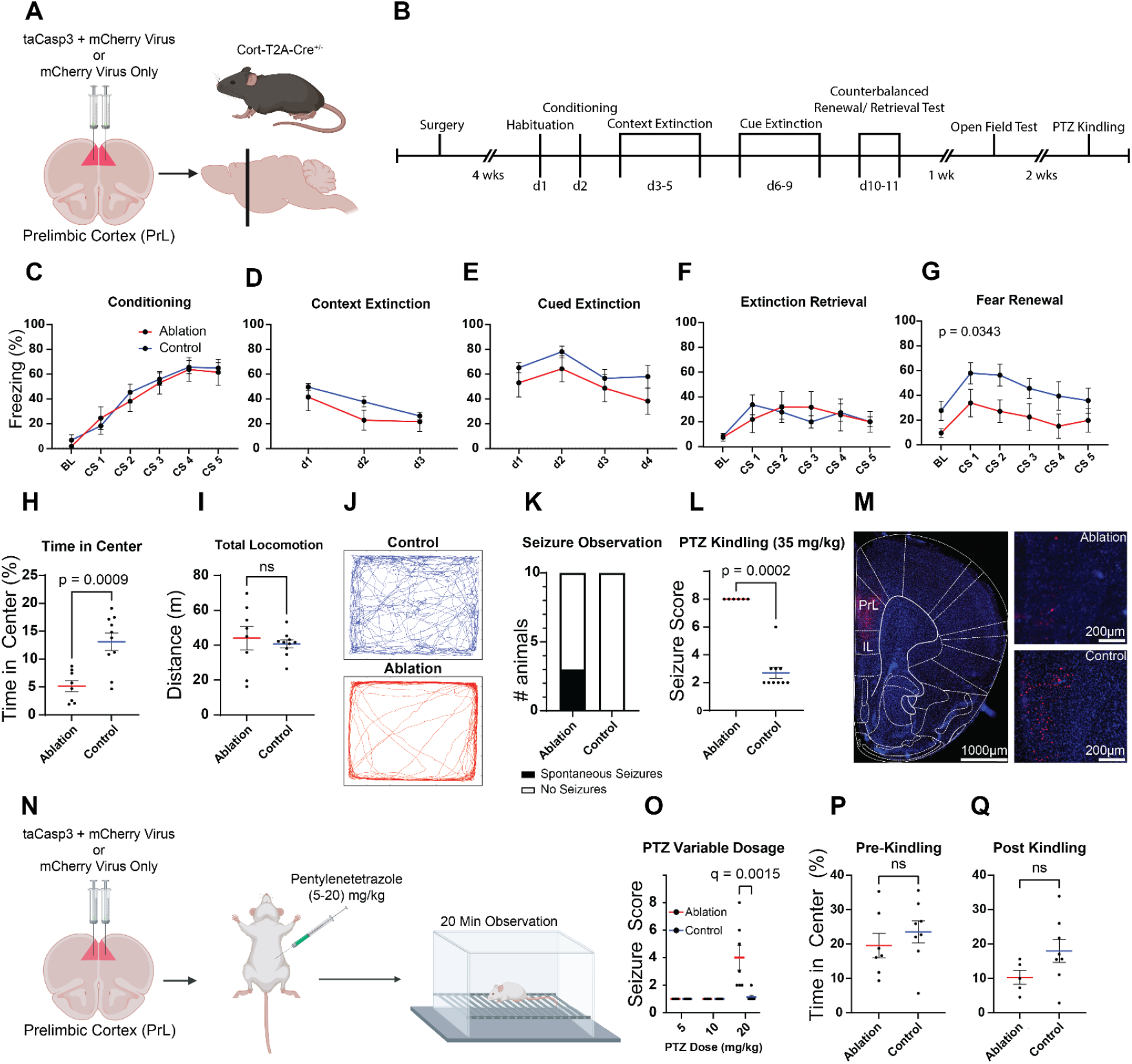
Ablation of cortistatin (CST+) neurons produces spontaneous seizures and increases sensitivity to pentylenetetrazole (PTZ) kindling. A) Schematic showing bilateral viral delivery of constructs encoding taCasp3/mCherry (Ablation group) or mCherry alone (Control group) into the prelimbic cortex (PrL) of *Cort*-T2A-Cre mice (n =10 per group, 5M and 5F) to ablate CST+ neurons. Created in BioRender. Lab, M. (2026) https://BioRender.com/fdzih8g B) Experimental timeline showing behavioral assays following surgical recovery and viral incubation, including auditory fear conditioning and extinction, open field testing, and PTZ kindling tests. C-G) Freezing behavior during fear conditioning (C), context extinction (D), cued extinction (E), extinction retrieval (F), and fear renewal (G), did not differ significantly between groups. H) In the open field, mice in the Ablation group spent less time in the center, but showed no change in total locomotion (I). J) Example traces of open field data from representative animals. K) During fear conditioning and extinction, a subset of mice in the Ablation group developed spontaneous, tonic-clonic seizures. L) Upon administration of a single sub-threshold dose (35 mg/kg) of PTZ, all mice in the Ablation group developed fatal seizures (Racine score of 8). M) Fluorescence microscopy images confirm selective loss of mCherry-labeled CST+ interneurons within PrL following ablation. N) Schematic illustrating PTZ kindling at increasing doses in a second cohort of mice (n = 8 per group; 4M and 4F). Created in BioRender. Lab, M. (2026) https://BioRender.com/q47vywk O) Seizure score increased in the Ablation group with increasing dose of PTZ. P-Q) Time spent in the center of the open field before (P) and after (Q) kindling shows no difference between groups. Data represent mean ± standard error of the mean (SEM).

To identify the acute dose threshold to induce seizures, we generated an additional cohort of Control and Ablation mice (n = 16; 8M, 8F). 4 weeks after surgery, these mice were administered daily, increasing doses of PTZ (5 - 20 mg/kg) until seizures developed in the Ablation group. Doses as low as 20 mg/kg were sufficient to induce seizures in the Ablation group (Fig. 1I, *U* = 6.000, *q* = 0.01078). In contrast to the first cohort, no differences between groups were observed in the open field before (*t* = 0.8371, *p* = 0.4176) or after (Fig. 1J, *t* = 1.680, *p* = 0.1211) PTZ kindling. Because the second cohort did not undergo fear conditioning, it is possible that the open field avoidance behavior observed in the first cohort (Fig. 1J) required interaction of CST+ ablation with a prior stressor.

### 2.2 Acute inhibition of PrL CST+ neurons increases seizure susceptibility in female mice

Our results established that ablation of PrL CST+ neurons induces spontaneous seizures and lowers the dose necessary to induce seizures with PTZ kindling. However, since ablation permanently impairs inhibitory tone, and loss of tonic inhibition can promote potentiation of excitatory synapses (C. Wu & Sun, 2015), (Tullis & Bayer, 2024), we investigated whether acute inhibition of CST+ neurons in PrL also impacted fear learning and seizure susceptibility. To test this, we bilaterally injected viral vectors encoding either a cre dependent hM4Di inhibitory designer receptor exclusively activated by designer drugs (iDREADD) (AAV8-Ef1a-DIO-hM4Di-mCherry; hM4Di group) or mCherry (AAV8-Ef1a-DIO-mCherry; Control group) into the PrL of *Cort*-T2A-cre^+/-^ mice (Fig. 2A). Four weeks after surgery, mice were tested in the auditory fear conditioning paradigm described above. Thirty min prior to cued extinction sessions, mice received an injection of clozapine-n-oxide (CNO, 5 mg/kg, i.p.), to activate the iDREADD and inhibit PrL CST+ neurons. No differences in measures related to fear learning or extinction were observed (Fig. S1). Following fear conditioning, mice underwent PTZ kindling. All mice were injected with CNO (5 mg/kg, i.p.) followed by PTZ (30 mg/kg, i.p.) 30 min later and were observed for 20 min for signs of seizures. Acute inhibition of PrL CST+ neurons in the hM4Di group increased the rate of seizure kindling compared to Control mice across days (Fig. 2C, *F*_(1,15)_ = 5.890, *p* = 0.0283, main effect of group). hM4Di mice took fewer days to fully kindle (defined as two consecutive sessions with Racine scale score ≥ 5) compared to Control mice (Figure 2D, *χ*^*2*^ = 4.277, *p* = 0.0378). This effect was driven entirely by female mice for seizure score (Fig. 2G, F_(1,9)_ = 12.97, *p* = 0.0057) and days to kindle (Fig. 2H, *χ*^*2*^ = 8.797, *p* = 0.0030). We saw no significant kindling observed in the male mice over the course of this kindling experiment (Fig. 2E, *F*_(1,9)_ = 2.770, *p* = 0.1471; Fig. 2F, *χ*^*2*^ = 0.000, *p* > 0.999). Last we observed no differences in locomotion (Fig. 2I, *t* = 0.3767, *p* = 0.7110) or time in center (Fig 2J, *t* = 0.7915, *p* = 0.4355) in the open field. Collectively, these data demonstrate that PrL CST+ neurons are critical to regulating cortical hyperexcitability.

**Fig. 2.**
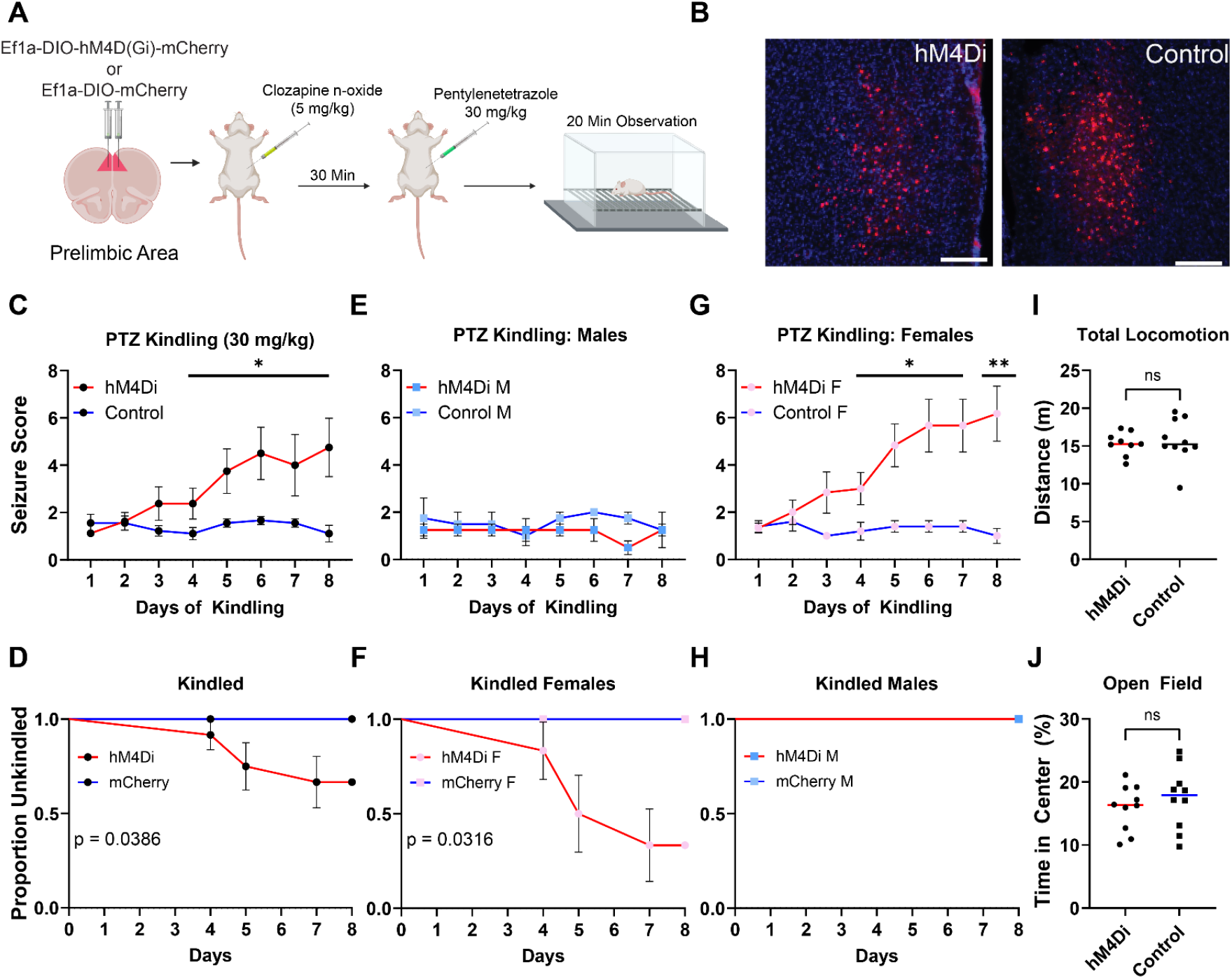
Inhibition of PrL CST+ neurons increases sensitivity to PTZ kindled seizures in female mice only. A) Experimental design showing bilateral injection of either a cre-dependent inhibitory DREADD (hM4Di group) or mCherry virus (Control group) into the PrL of *Cort*-t2A-Cre mice (n = 20, 10M and 10F). CNO (5 mg/kg) was administered 30 min prior to PTZ (30 mg/kg), and seizures were scored during a 20 min observation period. Created in BioRender. Lab, M. (2026) https://BioRender.com/mtj78g1 B) Representative fluorescence images showing robust expression of both viral constructs in the PrL. Scale bars: 200 μm C). Across 8 d of PTZ kindling, hM4Di mice exhibited greater seizure severity compared to Control mice. Sex-specific analyses further revealed that inhibiting CST+ neurons increased seizure susceptibility in female (G), but not male (E) mice. D, F, H) Survival analyses confirmed a sex-specific effect, showing that female mice in the hM4Di group take fewer days to kindle relative to controls. I,J) Behavioral tracking shows no differences in the open field between groups in total distance traveled (I) or time spent in the center (J). Data represent mean ± SEM. * = *p* > 0.05, ** = *p* > 0.01.

### 2.3 Kindled seizures broadly activate GABAergic subtypes in the PrL

After confirming that acute inhibition of CST+ neurons was sufficient to induce seizures, we investigated whether different subtypes of inhibitory neurons are preferentially impacted in response to kindled seizures. We generated a new cohort of Control and hM4Di (n = 5 per group, all female) as described above. Four weeks following surgery, mice in both groups were exposed to PTZ kindling. Mice in the hM4Di group were paired with mCherry expressing Control mice matched for age and sex. Once one animal of each pair reached a seizure score of 5 or higher, that animal and its paired counterpart were euthanized 2 h later, and brains were collected. We performed single molecule RNA fluorescent *in situ* hybridization (smFISH), to visualize *Fos* expression within inhibitory neuron subtypes marked by *Pvalb, Sst*, and *Cort* (Fig. 3A). We found significant recruitment of *Fos* in the hM4Di group following seizure compared to mCherry controls that did not experience seizures (Fig. 3B, *t* = 2.920, *p* = 0.0226). While we found no differences in proportions of *Pvalb*+ (*t* = 0.1502, *p* = 0.8844), *Sst*+ (*t* = 1.302, *p* = 0.2481), or *Cort*+ (*t* = 0.4878, *p* = 0.6395) neurons between either groups (Fig. 3B), all three subtypes of inhibitory neurons displayed elevated expression of *Fos* following seizures (*p*-values < 0.05) (Fig. 3C). ∼55% of *Cort*+ cells were also positive for *Sst* while ∼30% of *Cort*+ cells were positive for *Pvalb* (Fig. 3D), showing that while expressed within both these major inhibitory populations, CST+ neurons are primarily a population of Sst-expressing neurons. ∼50% of all *Sst*-expressing cells and 25% of *Pvalb*-expressing cells in the PrL coexpress *Cort* showing that CST+ neurons represent a distinct subset of these cell types (Fig. 3E). As expected, mice in the hM4Di group displayed increased seizure severity relative to mCherry Controls (Fig. 3F, *U* = 20, *p* = 0.0118).

**Fig. 3.**
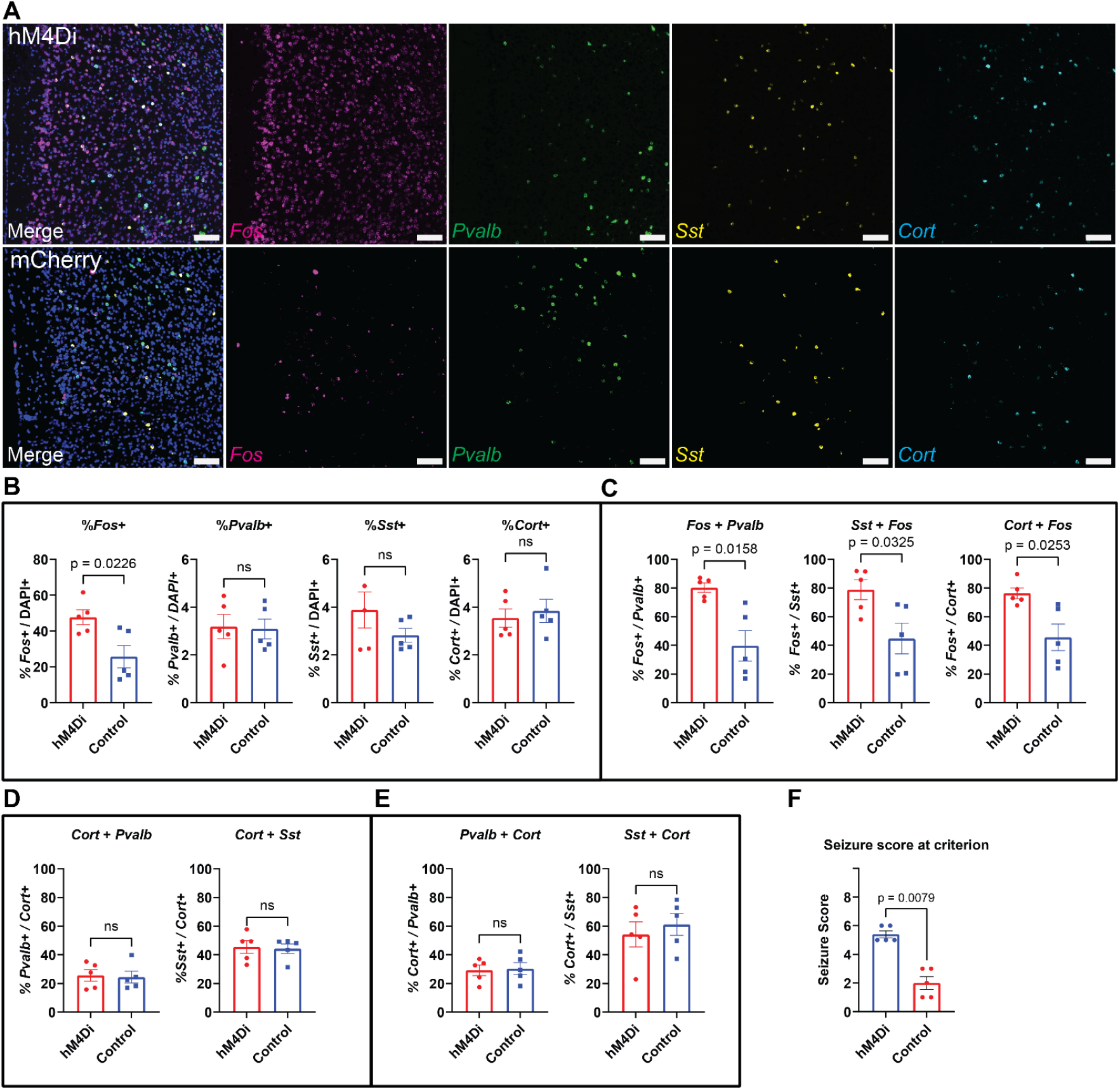
Recruitment of inhibitory neuron subtypes in response to inhibition of PrL CST+ neurons during PTZ kindling. A) Representative smFISH images from a hM4Di and mCherry control mice showing expression of *Fos* (magenta), *Pvalb (*green), *Sst* (yellow), and *Cort* (cyan) mRNA transcripts in the PrL following PTZ-kindled seizures. Scale bars: 200 μm. B) Quantification of total *Fos+, Pvalb*+, *Sst*+, and *Cort*+ cells (normalized to DAPI) revealed increased *Fos* expression following hM4Di-facilitated seizures. (n = 10 per group; 5F) C) Co-expression analyses revealed increased activation of *Pvalb*+, *Sst*+, and *Cort*+ neurons. D) *Cort*+ cells exhibited greater overlap with *Sst*+ (∼40%) relative to *Pvalb*+ (∼25%). These proportions do not differ between experimental groups. E) Similarly, *Sst*+ cells displayed higher proportions of *Cort* co-expression (∼50%) relative to *Pvalb*/*Cort* co-expression (∼25%) F) Seizure scores for hM4Di and mCherry control animals from the final day of kindling. Black data points indicate animals that were not used for smFISH. Data represent mean ± SEM. All p-values represent comparisons between groups performed using Welch’s t-test.

### 2.4 Inhibition of BDNF-TrkB signaling increases seizure susceptibility in female mice

Developmental deletion of TrkB in CST+ neurons causes spontaneous seizures that become fatal by 8 weeks of age (J. L. Hill et al., 2019), suggesting that BDNF-TrkB signaling is required for CST+ neurons to suppress hyperexcitability. However, whether BDNF-TrkB signaling is required to support normal CST+ neuron function in the adult brain is unknown. To address this question, we bilaterally injected either a cre dependent viral construct, encoding a truncated form of the TrkB receptor (TrkB dominant negative;TrkB.DN) or cre dependent mCherry (Control) viral construct into the PrL of adult *Cort*-t2A-cre^+/-^ mice (Fig. 4A, C). TrkB.DN lacks the catalytic tyrosine kinase domain that is necessary for downstream signaling but is still capable of binding BDNF (Fig. 4B). Therefore, expression of the mutant receptor reduces available BDNF to bind the native TrkB receptor, reducing BDNF-TrkB signaling in CST+ neurons specifically. 4 weeks after surgery, animals underwent PTZ kindling, with animals receiving a single dose of PTZ (35 mg/kg i.p.) daily followed by a 20 min observation window. We found no differences in mean seizure score across days between the TrkB.DN and Control groups (Fig. 4D), or when analyzing separately by sex (Fig. 4B-C). However, when we examined the latency to kindling (2 consecutive days of Racine score >=5), we found a significant effect of group (hazard ratio (HR):3.21, 95% confidence interval (CI) [1.05 - 9.91], *p* = 0.041), sex (HR: 4.78, 95% CI [1.59 - 14.37], *p* = 0.005), and a group x sex interaction (HR: 0.206, 95% CI: [0.054 - 0.866], *p* = 0.031) (Fig. 4E). When analyzed separately by sex, there was no significant effect of group in males (Fig. 4G, HR: 1.50, 95% CI [0.62 - 3.63], *p* = 0.366). Post hoc contrasts revealed that the TrkB.DN group kindled significantly faster than the Control group (Fig. 4I, HR: 0.310, 95% CI [0.101 - 0.952], *p* = 0.041). Last we saw no difference between groups in the open field before (Fig. 4J, *t* = 0.1532, p = 0.8797) or after kindling (Fig. 4K, *t* = 2.100, *p* = 0.0543).

**Fig. 4.**
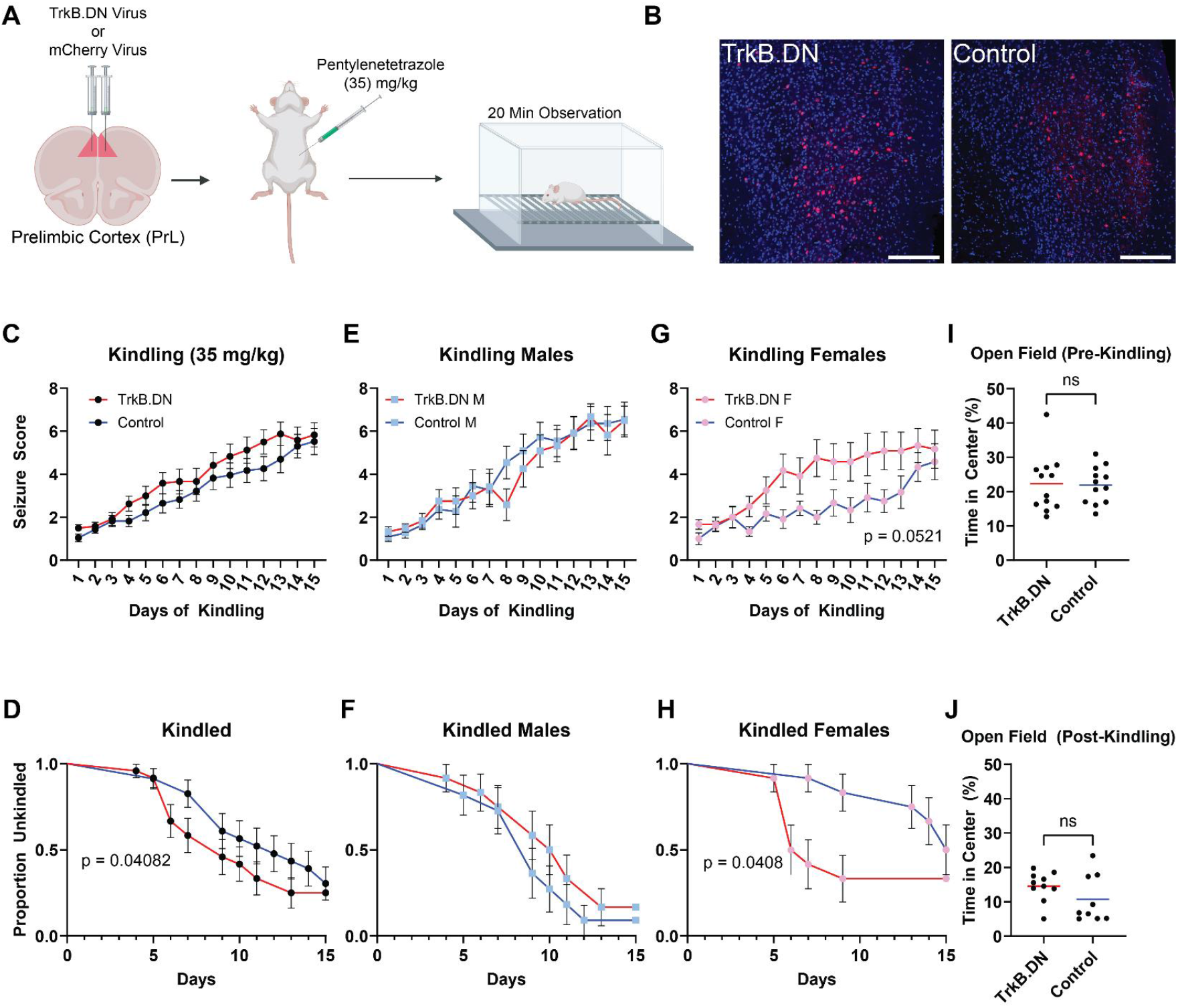
Disruption of TrkB signaling in PrL CST+ neurons increases seizure susceptibility only in female mice. A) Schematic illustrating viral injection of either cre-dependent dominant-negative TrkB (TrkB.DN) or mCherry (Control) vectors in the PrL of *Cort*-T2A-Cre mice, followed by daily administration of PTZ (35 mg/kg). Created in BioRender. Lab, M. (2026) https://BioRender.com/q47vywk B) Schematic showing native TrkB receptor and mCherry tagged dominant negative TrkB receptor, which lacks the intracellular catalytic domain. C) Immunohistochemistry showing robust viral expression in the PrL. D,F,H) Average seizure scores across 15 d of PTZ kindling showed no differences between groups in seizure score for the combined cohort or broken down by sex (n = 47; 24F, 23M). E,G,I) Survival analyses showing that female TrkB.DN mice take fewer days to kindle relative to controls. J,K) No differences in time spent in the center of an open field were observed before or after kindling. Data represent mean ± SEM. p-values in D,F,H represent main effect of group from two-way ANOVA; p-value E represents and main effect of group from cox proportional hazards model, and p-values from G,I represent contrast between effect of group for each sex.

## 3 Discussion

Despite evidence that epilepsy and stress-related psychiatric disorders are frequently comorbid (Chen et al., 2017; Pepi et al., 2024), little is known about the cellular mechanisms mediating shared etiology. Transcripts encoding marker genes for CST+ neurons are among among the most downregulated genes in postmortem human dACC tissue from individuals diagnosed with PTSD and MDD (Jaffe et al., 2022), suggesting CST+ neurons might be particularly impacted in these disorders. In addition, we found that constitutive ablation of CST+ neurons, or disruption of TrkB signaling in CST+ neurons, results in fatal spontaneous seizures in developing mice (J. L. Hill et al., 2019). However, whether disruption of CST+ neurons in adult mice would also impact seizure susceptibility or fear regulation remained unknown. We found that genetic ablation of CST+ neurons in the PrL of adult mice disrupted context-dependent fear renewal, resulted in spontaneous seizures, dramatically increased susceptibility to chemically induced seizures, and increased anxiety following stress. We additionally found that acute, daily inhibition of PrL CST+ neurons accelerates PTZ kindling exclusively in female mice, an effect that was recapitulated by disruption of BDNF–TrkB signaling in these neurons. Together, these findings identify CST+ neuron dysfunction as a potential common neurobiological mechanism linking stress-related psychiatric disorders and seizure vulnerability.

### PrL CST+ neuron dysfunction may represent a common mechanism explaining comorbidities between epilepsy and PTSD

Individuals with PTSD often display deficits in contextual processing and severe anxiety (Liberzon & Abelson, 2016; Maren et al., 2013), and our transcriptomic investigation of postmortem human brain tissue found that a number of genes commonly found in SST+ neurons (including *Cort*, which encodes the neuropeptide CST) were among the most strongly downregulated in the dACC of individuals with PTSD (Jaffe et al., 2022). Our investigation of the function of CST+ neurons in the adult rodent PrL has revealed that these neurons are largely a subpopulation of SST+ neurons and appear to be critical for context-dependent fear regulation. Specifically, genetic ablation of CST+ neurons in the PrL impaired context-dependent fear renewal of previously extinguished fear (Fig. 1G). Prior work found that inhibition of PrL SST+ neurons during auditory fear conditioning disrupted memory encoding and resulted in poor memory retrieval 24 hours later (Cummings & Clem, 2020). This study additionally showed that SST+ inhibition during retrieval alone was sufficient to disrupt memory retrieval (Cummings & Clem, 2020). Conversely, we found that genetic ablation of the CST+ subpopulation had no impact on fear encoding or retrieval and only impacted fear renewal. This suggests that CST+ neurons in the PrL are likely distinct from SST+ neurons that form part of the fear “engram” within the PrL (Cummings et al., 2022). This is thematically consistent with the fact that CST+ neurons commonly co-express a number of anxiolytic neuropeptides, including SST, CST, CRHBP, and NPY (Jaffe et al., 2022; Martinowich et al., 2011; Kristen R Maynard et al., 2020). Other work has shown that PrL SST+ neurons are critical for memory discrimination by demonstrating that inhibition of PrL SST+ neurons impairs rodents ability to discriminate between aversive and neutral stimuli, thus increasing fear generalization (Z. Wang et al., 2024). Although we did not explicitly test for fear generalization, we hypothesize that decrements in context discrimination in CST+ ablated mice might have resulted in blunted fear renewal observed in our first experiment (Fig.1G). That is, the inability of mice to appreciate a change in context prevented the return of previously extinguished fear. Finally, we observed a decrease in time spent in the open field (interpreted as an increase in anxiety-like behavior) following stress due to fear conditioning (Fig. 1H) in animals with ablated CST+ neurons, but not in animals that never underwent fear conditioning. Taking into account an extensive literature demonstrating the SST+ neurons are uniquely vulnerable to stress (Girgenti et al., 2019; Lin & Sibille, 2015; Tomoda et al., 2022), we speculate that decrements in CST+/SST+ neuron function might be a critical feature underlying increased susceptibility to stress and stress-related psychiatric symptoms. Determining the precise role and function of PrL CST+ subpopulations of SST+ neurons in fear memory processes, and how this function is impacted by stress, should be a clear line of investigation for future studies.

### CST+ neurons are critical for maintaining E/I balance in adult PrL

Our findings demonstrate that loss of CST+ neuronal function leads to increased seizure vulnerability. Previous work found that intracerebroventricular administration of CST attenuates seizure severity and protects against cell death in chemoconvulsant seizure models through its action on SST receptors (Braun et al., 1998; Aourz et al., 2014), and our lab previously demonstrated that constitutive ablation of CST+ neurons leads to spontaneous fatal seizures in early postnatal development (J. L. Hill et al., 2019). However, it was unknown whether seizures occurred directly due to the loss of inhibition provided by CST+ neurons or from aberrant development of cortical microcircuitry due to loss of CST+ neuron signaling. *Cort* expression rapidly increases in the second week of mouse postnatal development, reaching a peak at postnatal day 15 (de Lecea et al., 1997), a timepoint corresponding to maturation of the cortical inhibitory system (Okaty et al., 2009). Inhibitory signaling is essential for development of cortical networks (Ganguly et al., 2001; Hensch & Stryker, 2004; Che et al., 2018), so constitutive loss of CST+ neurons could induce seizures through developmental mechanisms or through disruption of E/I balance by loss of inhibitory tone. Here, we show that loss of CST+ neurons in adult mice is sufficient to induce spontaneous seizures and increase sensitivity to the kindling effects of PTZ (Fig. 1H-Q). Perhaps most striking is that we observed this phenotype following ablation of CST+ neurons exclusively within the PrL. This supports the idea that these neurons are critical for maintaining E/I balance in adult animals, and observed seizures are not due to abnormalities in inhibitory circuit development.

While the hippocampus and basolateral amygdala are well-studied as seizure foci, we also demonstrate importance of the PrL in the development and propagation of seizures. This is in line with our work showing that constitutive knockout of TrkB in CST neurons produces a similar seizure phenotype, and that these seizures were observed first in the cortex and then spread to the hippocampus (J. L. Hill et al., 2019). EEG recordings of both spontaneous generalized seizures and PTZ-induced seizures show initial activity in the prefrontal cortex that spreads to thalamus (Goldring, 1972). Further, recurrent circuits between PrL (and other cortical regions) and thalamus are implicated in the propagation of generalized seizures from seizure initiation sites (Meeren et al., 2002; Brodovskaya & Kapur, 2019; Niemeyer et al., 2022; Wicker et al., 2022). Taken together, this evidence supports a role for the PrL as a key region in the initiation and propagation of seizures and that loss of inhibition from CST neurons in the PrL can exacerbate the effects of seizure promoting stimuli.

### Sex differences in BDNF-TrkB confer differential susceptibility to kindled seizures after loss of CST+ neuron function

While localized ablation of CST+ neurons in adult mice was sufficient to drive spontaneous seizures and increase sensitivity to PTZ (Fig. 1K-L), this effect could arise from loss of inhibition provided byCST+ neurons at the time of PTZ administration or due to changes in plasticity downstream of chronic loss of tonic inhibition. Because tonic inhibition impairs potentiation of excitatory synapses (Z. Wu et al., 2014), its loss can strengthen excitatory connections within a circuit. To investigate whether acute loss of inhibition provided by CST+ neurons could exacerbate PTZ kindled seizures similarly to ablation, we acutely inhibited CST+ neurons in the PrL. Acute inhibition only affected sensitivity to PTZ in female mice (Fig. 2C-H), which is in line with literature demonstrating sex differences in SST+ neuron function. In mice, PrL SST+ neurons exhibit sex-specific transcriptional profiles that parallel stress-induced transcriptomic alterations following chronic stress (Girgenti et al., 2019). Further, disinhibition of SST+ neurons in mPFC promotes stress resilience in male, but not female mice (Jefferson et al., 2020), suggesting functional differences in SST+ populations across sex. Given these known sex differences in SST function, and the fact that a large portion of CST+ neurons in the PrL co-express SST (Fig. 3E), we speculate that a SST+ subset of CST+ neurons likely exhibits similar sexually divergent responses to PTZ or other forms of excessive neural excitation. This ultimately results in increased seizure susceptibility in SST+ microcircuits and grants differential vulnerability to excessive excitation as has been observed previously in stress models (Jiang et al., 2025).

Another possible explanation for these observations is that sex differences in BDNF-TrkB signaling in CST+ neurons account for the observed sex differences. We previously demonstrated that constitutive knockout of *Ntrk2* (the gene encoding TrkB) in CST+ neurons causes spontaneous seizures (J. L. Hill et al., 2019). Additionally, BDNF-TrkB signaling regulates expression levels of many genes encoding neuropeptides such as NPY, SST, CST, and CRHBP (Glorioso et al., 2006; Martinowich et al., 2011; Adachi et al., 2018), all of which are expressed within CST+ neurons. We also showed that knockout of BDNF-TrkB signaling in CST+ neurons alters expression of *Npy, Sst*, and *Cort* within these neurons (K R Maynard et al., 2021). Here, we show that antagonizing TrkB signaling in CST+ neurons in PrL reproduces the sex effect observed after chemogenetic inhibition (Fig. 4 D-I). Female mice have lower levels of BDNF expression in the cortex compared to males (Szapacs et al., 2004), so they may be more sensitive to perturbations that disrupt BDNF-TrkB signaling. Supporting this interpretation, BDNF haploinsufficiency increases levels of phosphorylated TrkB in the frontal cortex of male mice, but reduces levels in females (R. A. Hill & van den Buuse, 2011). As TrkB phosphorylation is necessary to engage downstream BDNF-TrkB signaling, we can infer that signaling may be higher in males and lower in females in response to TrkB knockdown. As such, males may compensate for reduced bioavailable BDNF by increasing TrkB phosphorylation, while females do not. This would explain why only females show increased sensitivity to PTZ kindled seizures following knockdown of TrkB signaling. As BDNF regulates neuropeptides that promote inhibition in CST+ neurons, and BDNF itself promotes neural activity (Bolton et al., 2000), loss of TrkB signaling in female mice may weaken the ability of CST+ neurons to respond homeostatically to increased neural activity. Last, given known sex differences in epilepsy prevalence and presentation (Reddy et al., 2021), CST+ neuron dysfunction may account for some of these differences. For instance, idiopathic generalized epilepsies are more commonly observed in women than men (Christensen et al., 2005), and women are more likely to have treatment resistant epilepsy (Cepeda et al., 2022).

### Conclusions

In summary, we establish that loss of CST+ neurons in the PrL impairs context-dependent fear renewal and induces anxiety only after stress exposure. We show that loss of CST+ neurons in the PrL of adults can induce seizures. We establish that loss of function in CST+ neurons leads to increased vulnerability in females to PTZ induced seizures, implicating CST+ neurons as a cellular population that may explain sex differences in epilepsy. We also show that this population can regulate phenotypes relevant for stress-related psychiatric disorders as well as susceptibility to seizure, supporting evidence for common pathology between these disorders that may explain observed comorbidities in humans.

## 2 Methods

### 4.1 Animals

Heterozygous male and female *Cort*^*tm1(cre)Zjh*^/J (referenced in text as CST+^Cre^, Strain #:010910, Jackson Laboratory, Bar Harbor, ME) were used for all experiments. These mice express Cre-recombinase under the control of the endogenous *Cort* promoter. Mice were group housed (2-5 per cage) in disposable polycarbonate caging (Innovive, San Diego, CA). Cages were housed in a temperature and humidity-controlled environment with a reverse 12/12 light/dark cycle (lights on at 19:00 hours / lights off at 07:00 hours). Mice received water and Teklad Irradiated Global 16% Protein Rodent Diet (#2916; Envigo, Indianapolis, IN) in the home cage ad libitum for the duration of the experiments. Behavioral testing was conducted Monday-Friday during the dark phase (07:00–19:00 hours). All experiments and procedures were approved by the Johns Hopkins Animal Care and Use Committee and in accordance with the Guide for the Care and Use of Laboratory Animals.

### 4.2 Surgical Procedures

Mice were anesthetized with isoflurane (induction: 2–4% in oxygen, maintenance: 1–2%) and secured to a stereotaxic frame. Incision sites were disinfected with 10% betadine solution, local anaesthetic (2% lidocaine) was injected under the scalp, and analgesic (20 mg/kg meloxicam) was injected intraperitoneally (IP) before any incisions were made. The skull was exposed by incision along the midline of the scalp, and the skull was leveled using bregma and lambda as reference points. A small hole was drilled with a 0.9 mm burr (Fine Science Tools, Foster City, CA) above the PrL, and 350 nl of a viral vector was injected into the PrL (AP: +1.75; ML: ±0.3.5, DV: −2.1). For ablation experiments mice received either an injection of AAV5-Ef1a-flex-taCasp3 (Addgene, #45580-AAV5) combined with AAV5-Ef1a-flex-mCherry at a 1:9 ratio, or just the AAV5-Ef1a-DIO-mCherry (UNC Vector Core) as a control. For inhibitory DREADD experiments, animals received either AAV9-hSyn-DIO-hM4D(G_i_)-mCherry (Addgene, #44362) or AAV9-hSyn-DIO-mCherry (Addgene, #50459-AAV9) as a control vector. For the TrkB knockdown experiments, mice received either AAV5-Ef1a-DIO-TrkB.DN-mCherry (UNC Vector Core) or AAV5-Ef1a-DIO-mCherry (UNC Vector Core) as a control vector. Injections were made using a Micro4 controller and UltraMicroPump along with a 10 μl Nanofil syringes equipped with 33-gauge needles (WPI Inc., Sarasota, FL). The syringe was left in place for 5 minutes after injection to minimize diffusion. After injections were completed, the incision was closed using surgical staples (Fine Science Tools, Foster City, CA), and animals recovered on a heating pad for 30–60 mins before being returned to the colony room. Animals were closely monitored for health and recovery progress and received Meloxicam injections (20 mg/kg) to relieve pain for three additional days.

### 4.3 Behavioral Assays

#### 4.3.1 Auditory Fear Conditioning

All days of the cued auditory fear conditioning task were performed inside sound attenuated operant conditioning chambers (Clever systems) equipped with chamber lights containing a cage equipped with cage lights, audio speaker, and electrifiable parallel bar grid floor. Chambers were modified as needed to achieve different contexts throughout the experiments. On the days of behavior animals were transferred from the colony room in plastic transport boxes to the behavioral rooms. The chamber Contexts are as follows: **Context A:** Cages were cleaned with 1% ammonia, white room lights were turned on, chamber and cage lights were left off, chamber doors were left open, parallel bar grid floor was used, and no bedding was used in the transport box. **Context B**: Cages were cleaned with 70% ethanol, red room lights were turned on, chamber lights were turned off, cage lights were turned on, chamber doors were closed, black acrylic floor was placed over parallel bar flooring, and bedding was used in the transport box. **Context C:** Caged were cleaned with 1% virkon^®^ (Lanxess), red room lights were turned on, chamber and cage lights were turned on, chamber doors were left open, black acrylic floor was placed over parallel bar flooring, and no bedding was used in the transport box.

##### Context Habituation

Before undergoing conditioning, animals were habituated to each context on consecutive days. The animals were placed in the chamber for 20 minutes, and an overhead camera recorded their movement through the chamber. Freezing was quantified using automated tracking software (FreezeScan, Clever Systems inc). This software detects movement based on changes in pixel intensity between frames in the video recording, and if no movement is detected for at least 0.5s, the software records that as a period of freezing, and records the duration of detected freezing.

##### Auditory Cued Conditioning

Mice were placed into context A and following a 180 second baseline period a total of 5 tones were presented (10 sec, 75 dB, 3000 hz) followed by a footshock (1 sec, 0.75 mA) at tone offset with a 60 second intertrial interval between tone onsets. Freezing was quantified at baseline and for the 60 second interval following each tone onset.

##### Cued Extinction

Mice were placed into context B and following a 180 second baseline period a total of 30 tones were presented (10 sec, 75 dB, 3000 hz) in the absence of foot shocks with a 30 second interval between tone onsets. Freezing was quantified at baseline, and across blocks of 3 tone presentation. Cued extinction sessions continued across days until animals stopped showing elevated freezing to the tone. For experiments with hM4D(G_i_) animals, clozapine n-oxide (CNO) was administered (5 mg/kg, IP) to both hM4D(g_i_) and mCherry control animals 30 minutes prior to cued extinction sessions.

##### Context Extinction

Mice were placed into context A and observed for 20 minutes with no tone or footshock presentations. Freezing was quantified using automated tracking software. Context extinction was repeated across days until the time spent freezing to the context plateaued.

##### Cued Fear Retrieval and Fear Renewal

Mice were sorted randomly into two groups and were tested in either the cued extinction context (context B) or a novel context (context C) for fear retrieval and renewal respectively. In both contexts, following a 180 second baseline period, 5 tones were presented (10 sec, 75 dB, 3000 hz) with a 60 second interval between tone onsets. Freezing was quantified at baseline and for the 60 second interval following each tone onset. One day later mice were placed in the opposite context of the first day and tested again for fear retrieval or fear renewal.

#### 4.3.2 Pentylenetetrazole Kindling

For all kindling experiments, mice repeatedly receieved IP injections of pentylenetetrazole (PTZ) (35 mg/kg, unless otherwise noted) of and were immediately placed into an empty cage without bedding for a 20 minute observation period. For caspase ablation experiments, mice were started at a dose of 10 mg/kg that was ramped up to 20 mg/kg across several days. For the hM4D(G_i_) experiments, mice received dosages of 30 mg/kg each day. For TrkB dominant negative experiments, mice received doses of 35 mg/kg each day. For experiments involving hM4D(G_i_), all mice were given a dose of CNO (5 mg/kg, IP) 30 minutes before the dose of PTZ. Mice were monitored for signs of seizure and seizure severity was scored on a modified Racine seizure severity scale (Li et al., 2014). Scores for this scale are as follows: 1) Animal freezes in place for at least 5 seconds. 2) Animal jerks with ears going flat. 3) Animal jerks accompanied with tail flicking upward > 90°. 4) forelimbs extend and push the body upward in a tonic state. 5) generalized clonic seizure with loss of righting ability. 6) Mouse begins running and jumping. 7) Running and jumping ceases and the animal extends hindlimbs beyond their body. 8)Arrest of breathing and death. Latency from injection to each seizure severity stage was also recorded. Animals received doses of PTZ 5 days a week, with saturday and sunday off in between, and PTZ administration was continued until animals reached their experimental endpoint for seizure severity. For all experiments, this was until each animal reached a seizure severity score of 5.

#### 4.3.3 Open Field

Animals were placed into a 60cm x 60cm square open field chamber, and were observed for 10 minutes. Position data for open field testing were captured with CaptureStar software (CleverSys, Inc.), and occupancy maps, time spent in the center, and number of center crossings were analyzed in software.

### 4.4 Immunohistochemistry

Mice were anaesthetized with 2% isoflurane and euthanized by transcardial perfusion with 4% formaldehyde. Brains were post-fixed for at least 12 hours, and cryo-protected by soaking in a 30% sucrose solution. Serial sections encompassing the PrL were cut using a microtome. (Leica Biosystems Inc., Wetzlar, Germany) equipped with a freezing stage (Physitemp, Cliffton, NJ). Sections were incubated with anti-mCherry. Sections were washed with 1x phosphate buffered saline (PBS) for 10 minutes, followed by a 20-minute incubation in PBS + 0.5% Triton x-100 (PBST) to permeabilize the sections. Sections were then blocked in 10% normal goat serum in 0.1% PBST for 1 hour at room temperature. Following blocking, sections were incubated with a rabbit anti-mCherry antibody (Invitrogen, cat# PA5-34974) at a 1:1000 dilution in blocking buffer overnight at 4 °C. After primary incubation, sections were washed 3x with 0.1% PBST for 10 minutes. Secondary antibody consisting of a goat anti-rabbit antibody conjugated to alexafluor-555 (Invitrogen, Cat# A-21428) was then applied for 2 hours at room temperature at a 1:1000 dilution. During the last 30 minutes of incubation DAPI solution was added at a 1:30000 dilution. Sections were washed 3x times with 0.1% PBST for 10 minutes. Sections were then mounted onto superfrost microscope slides (Fisher Scientific), fluoromount G mounting medium was applied, and a coverslip was affixed over the sections. Slides were then imaged on a confocal fluorescent microscope to confirm localization and expression of viruses.

### 4.5 Single molecule fluorescence *in situ* hybridization

Mice were euthanized by cervical dislocation, and brains were dissected from the skull. Brains were immediately flash frozen in chilled isopentane. Brains were equilibrated to -15°C for at least 1 h prior to cryosectioning (Leica Biosystems). 15 *μ*m serial cryosections through PrL were collected, mounted onto Superfrost microscope slides (Fisher Scientific) and stored at -80°C, until experimental use. Single molecule RNA fluorescent *in situ* hybridization assays were performed using the RNAscope fluorescent multiplex kit v3 according to manufacturer’s instructions as previously described (K R Maynard et al., 2021). Briefly, tissue sections were fixed with a 10% neutral buffered formalin solution for 30 min at room temperature, series dehydrated in ethanol, and pretreated with hydrogen peroxide at RT for 10 min then with protease IV for 30 min. Sections were incubated with probes against *Cort, Pvalb, Fos*, and *Sst* (Cat #s 421931,404631-C2, 490761-C3, 316921-C4) for 2 h and stored overnight in 4x SSC (saline-sodium citrate) buffer. Probes were fluorescently labeled with Opal dyes (PerkinElmer, Waltham, MA). Opal dyes were diluted at 1:500 and assigned to each probe as follows: Opal520 to *Fos*, Opal570 to *Cort*, Opal620 to *Pvalb*, and Opal690 to *Sst*. Tiled confocal fluorescent microscope images were captured on a Nikon eclipse Ti-2 microscope equipped with a 20x/1.4A objective. Linear unmixing was performed as previously described (Ramnauth et al., 2025), and images were processed using Halo analysis software (Indica Labs).

### 4.6 Statistical Analyses

Statistics were calculated using GraphPad Prism Software (GraphPad Software, La Jolla, CA). For all animal studies, the reported *N* represents the total number of animals in the study. These N are reported in both the Methods section as well as in the individual figure legends for the respective experiments. Data comparing means between two groups was compared using an unpaired student’s t-test for parametric data including open field test data and cell counting. To compare means between two groups with non-parametric data, Wilcoxon signed-rank test, including assays where seizure score was a dependent variable. To compare group differences across timepoints and sex, two-way repeated measure analysis of variance (ANOVA) was performed. For survival analyses, Kaplan-Meier curves were analyzed by the Cox proportional hazards analysis. All alphas were set at 0.05.

## 5 Acknowledgements, Funding, Authorship Contributions

Acknowledgements

We thank Suhaas Adiraju, Jorge Miranda-Barrientos, Jason Rehg, and Robert Phillips for technical assistance with animal experiments. We also thank Haya Algrain and Stephanie Cerceo Page for their training and assistance in imaging and analyzing smFISH data within the LIBD Microscopy and Spatial Biology Core. Portions of some figures were created with BioRender.com.

## Author Contributions

Conceptualization: KM, MST

Formal Analysis: AJS, MST

Investigation: AJS, LF, MST

Data Curation: AJS, MST

Writing – original draft: AJS

Writing – review & editing: AJS, MST, KM

Supervision: KM, MST

Funding acquisition: KM, MST

Project Administration: KM, MST

## 8 Supplementary Figures

**Fig. S1.**
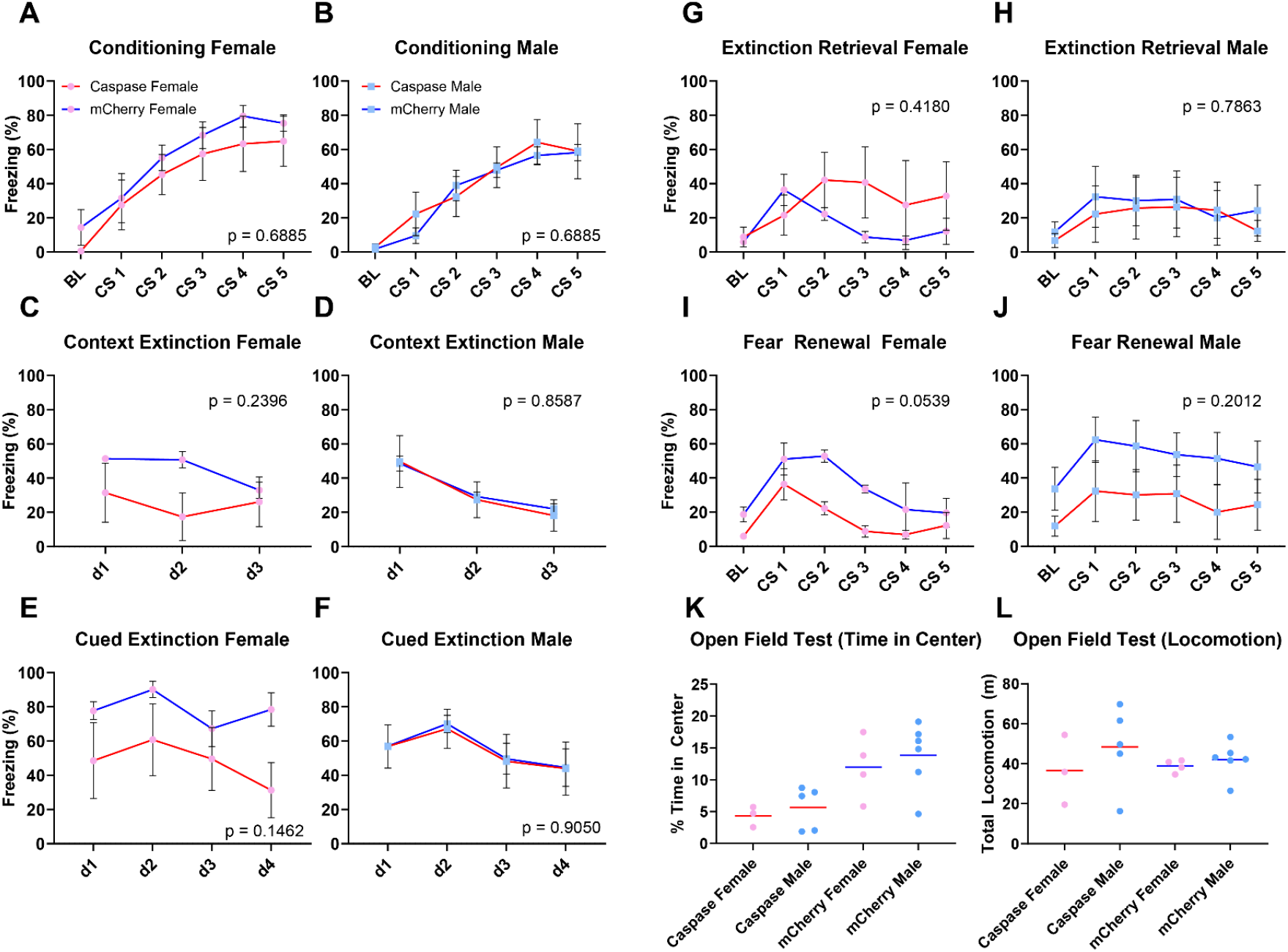
Cst+ neuron ablation shows no sex differences during fear conditioning. A-L) When broken down by sex, there were no observed differences in any phase of the fear conditioning paradigm including during conditioning (A-B), context extinction (C-D), Cued Extinction (E-F), fear extinction retrieval (G-H),or fear renewal (I-J). (K-L) There were no observed differences between groups or sex in open field in time spent in center (K) or total locomotion (L).

**Fig. S2.**
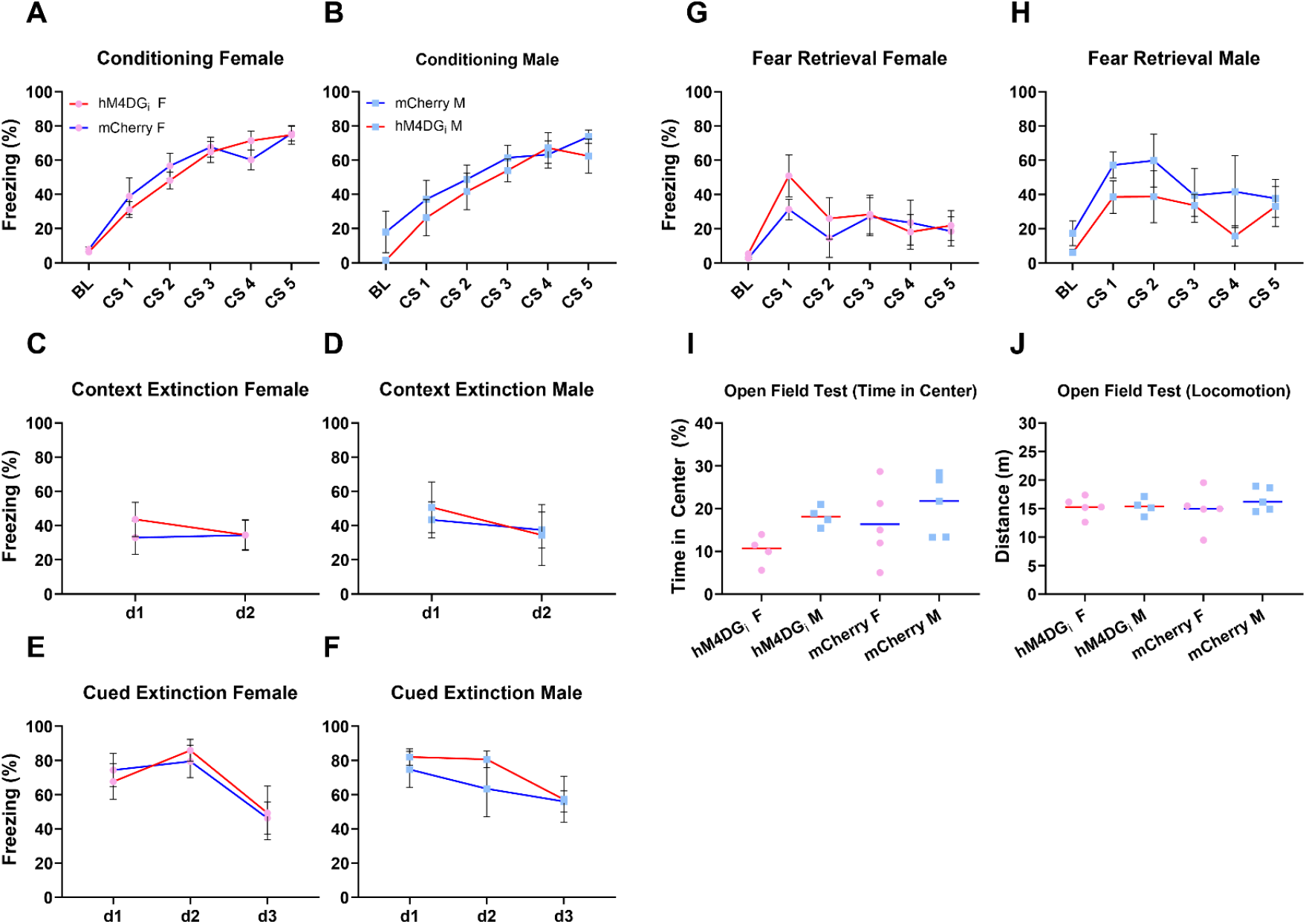
DREADD inhibition of Cst+ neurons during cued extinction has no effect on extinction learning. A-B) Both males and females condition normally to footshock-tone presentations. C-H) No differences are observed in contextual fear extinction (C-D), cued fear extinction (E-F), or extinction retrieval (G-H). (I-J) No significant differences were observed in open field testing in time spent in center (I) or total locomotion (J)

